# Structure-based prediction of T cell receptor:peptide-MHC interactions

**DOI:** 10.1101/2022.08.05.503004

**Authors:** Philip Bradley

## Abstract

The regulatory and effector functions of T cells are initiated by the binding of their cell-surface T cell receptor (TCR) to peptides presented by major histocompatibility complex (MHC) proteins on other cells. The specificity of TCR:peptide-MHC interactions thus underlies nearly all adaptive immune responses. Despite intense interest, generalizable predictive models of TCR:peptide-MHC specificity remain out of reach; two key barriers are the diversity of TCR recognition modes and the paucity of training data. Inspired by recent breakthroughs in protein structure prediction achieved by deep neural networks, we evaluated structural modeling as a potential avenue for prediction of TCR epitope specificity. We show that a specialized version of the neural network predictor AlphaFold can generate models of TCR:peptide-MHC interactions that can be used to discriminate correct from incorrect peptide epitopes with substantial accuracy. Although much work remains to be done for these predictions to have widespread practical utility, we are optimistic that deep learning-based structural modeling represents a path to generalizable prediction of TCR:peptide-MHC interaction specificity.

## Introduction

The specificity of T cell receptors (TCR) for peptides presented by major histocompatibility complex proteins (pMHC) is a critical determinant of adaptive immune responses to pathogens and tumors and of autoimmune disease. A predictive model of TCR:pMHC interactions, capable of mapping between TCR sequences and pMHC targets, could lead to advances in cancer immunotherapy and in the diagnosis and treatment of infectious and autoimmune diseases. Despite recent progress in TCR sequence analysis and modeling (Gielis et al., 2019; Huang et al., 2020; Mayer-Blackwell et al., 2021; Winther et al., 2021), a generalizable predictive model of TCR:pMHC interactions remains out of reach: existing predictors can learn to recognize new TCR sequences specific for pMHCs in their training set, but robust generalization to unseen pMHC epitopes has not been convincingly demonstrated (Moris et al., 2021). Two key difficulties are the diversity of TCR:pMHC recognition modes, a consequence of TCR sequence and structural diversity and flexibility in TCR:pMHC docking orientation, and the limited number of experimentally validated TCR:pMHC interaction examples for use in training.

We hypothesized that 3D structural modeling might offer a path toward generalizable prediction of TCR:pMHC interactions in the current data-limited regime. At the biophysical level, TCR:pMHC interaction specificity is determined by the structures and flexibilities of the interacting partners. A wealth of structural studies have provided valuable insights into the atomistic determinants of specificity (Rossjohn et al., 2015; Rudolph et al., 2006; Singh et al., 2017). Collectively, these experimentally determined structures define a range of docking geometries that likely covers the majority of unseen interactions; they also provide valuable templates for cutting-edge deep neural network structure prediction methods such as AlphaFold (Jumper et al., 2021) and RoseTTAfold (Baek et al., 2021). These prediction tools feature advanced network architectures with millions of parameters that are trained on structurally characterized proteins and their sequence homologs. Despite being trained on monomeric structures, these approaches can generate state-of-the-art structure predictions for protein complexes, and they have even been used to predict whether or not protein pairs will associate (Humphreys et al., 2021).

Here we show that a version of AlphaFold specialized for TCR:pMHC modeling can be used to predict TCR:pMHC binding specificity with some success. Whereas the default AlphaFold version trained to predict protein:protein docking (AlphaFold-Multimer (Evans et al., 2021)) shows inconsistent performance on TCR:pMHC structures, our specialized pipeline demonstrates improved accuracy and reduced computational cost. Moreover, this modeling pipeline has significant power to discriminate target peptides from decoy peptides as evaluated on a benchmark of human and mouse MHC class I epitopes. Importantly, success in predicting the correct peptide target correlates with structural accuracy of the models, suggesting that when the pipeline succeeds, it does so by recapitulating key specificity determinants. This work, together with previous studies applying molecular modeling techniques to TCRs (Borrman et al., 2020; Jensen et al., 2019; Lanzarotti et al., 2018; Pierce & Weng, 2013), suggests that structure-based approaches represent a promising path forward for predicting TCR:pMHC interaction specificity.

## Results

### Structure Prediction

We first evaluated the structure prediction performance of a recently released version of AlphaFold (AlphaFold-Multimer (Evans et al., 2021)) that was specifically trained for protein:protein docking. AlphaFold-Multimer leverages inter-chain residue covariation observed in orthologs of the target proteins to identify amino acid pairs making interface contacts. Given that TCR:pMHC interactions are determined in part by highly variable, non-germline encoded CDR3 regions, it was unclear whether AlphaFold’s strong docking performance on other systems would translate to TCR:pMHC interactions. Indeed, the AlphaFold-Multimer developers noted that it does not perform well on antibody:antigen complexes, which share many features with TCR:pMHC complexes.

We tested two versions of AlphaFold-Multimer, one in which the full sequences of the interacting partners are provided as input (‘AFM_full’: MHC-I or MHC-IIa, beta-2 microglobulin or MHC-IIb, peptide, TCRa and TCRb variable and constant domains), and one in which only the directly interacting domains are provided as input (‘AFM_trim’: TCR constant domains, beta-2 microglobulin, and C-terminal MHC domains are removed). Restricting to the core interacting domains speeds the calculations substantially at the risk of introducing decoy docking sites at the location of interfaces with the missing domains. Although both models were capable of generating high quality predictions on a nonredundant set of 130 TCR:pMHC complexes (as indicated by CDR loop RMSDs at and below ~2Å; details below), prediction quality was highly variable, and visual inspection revealed that many of the predicted models had displace peptides and/or TCR:pMHC docking modes that were outside the range observed in native proteins (**Fig. S1**). Additionally, these AlphaFold predictions took multiple hours per target to complete, limiting their throughput.

One limitation of AlphaFold-Multimer is that it does not support multi-chain templates (Evans et al.,2021): template information from the database of solved structures can inform the internal conformation of individual chains, but it does not guide the docking of chains into higher order complexes. The constrained nature of the TCR:pMHC binding mode suggests that higher and more consistent prediction accuracy could be obtained by providing additional template information. A challenge when modeling TCR structures is that the V-alpha and V-beta genes largely determine the best structural template, and these genes associate freely rather than in fixed pairings, which means that the optimal structural template for the TCR-alpha chain will often come from a different PDB structure as that for the TCR-beta chain. Additionally, the TCR:pMHC docking mode varies widely within an overall diagonal binding mode, in a way that is not easily predicted directly from sequence, making it challenging to select an optimal template for the TCR:pMHC relative orientation. Guided by these considerations, we developed an AlphaFold-based TCR docking pipeline that uses hybrid structural templates to provide a broad, native-like sampling of potential docking modes (**Fig. 1**). In this approach, individual chain templates are first selected based on sequence similarity to the target TCR:pMHC (**Fig 1A**). Hybrid complexes are created from these individual chain templates by using a diverse set of representative docking geometries to orient the TCR chains relative to the pMHC (see Methods). Docking geometries are defined in terms of the 6 degrees of freedom that relate the MHC reference frame to the TCR reference frame, where the MHC and TCR reference frames are defined based on internal pseudosymmetry (**Fig. 1B,D** and Methods). These hybrid complexes are provided as templates to multiple independent AlphaFold simulations, four templates per simulation, with the highest confidence model from the simulations taken as the final prediction (**Fig. 1C**). During benchmarking, templates and docking geometries from structures with similar TCRs or pMHCs to the target are excluded to reduce bias toward the native structure (see Methods; this constraint was not applied to the default AlphaFold-Multimer methods). Given that we are providing template information that constrains the inter-chain docking, we chose not to include additional multiple sequence alignment (MSA) information beyond the target sequence. This greatly speeds the predictions: MSA building is the most time-consuming part of the AlphaFold pipeline, and the neural network inference step is also significantly faster without MSA information.

**Figure 1.**
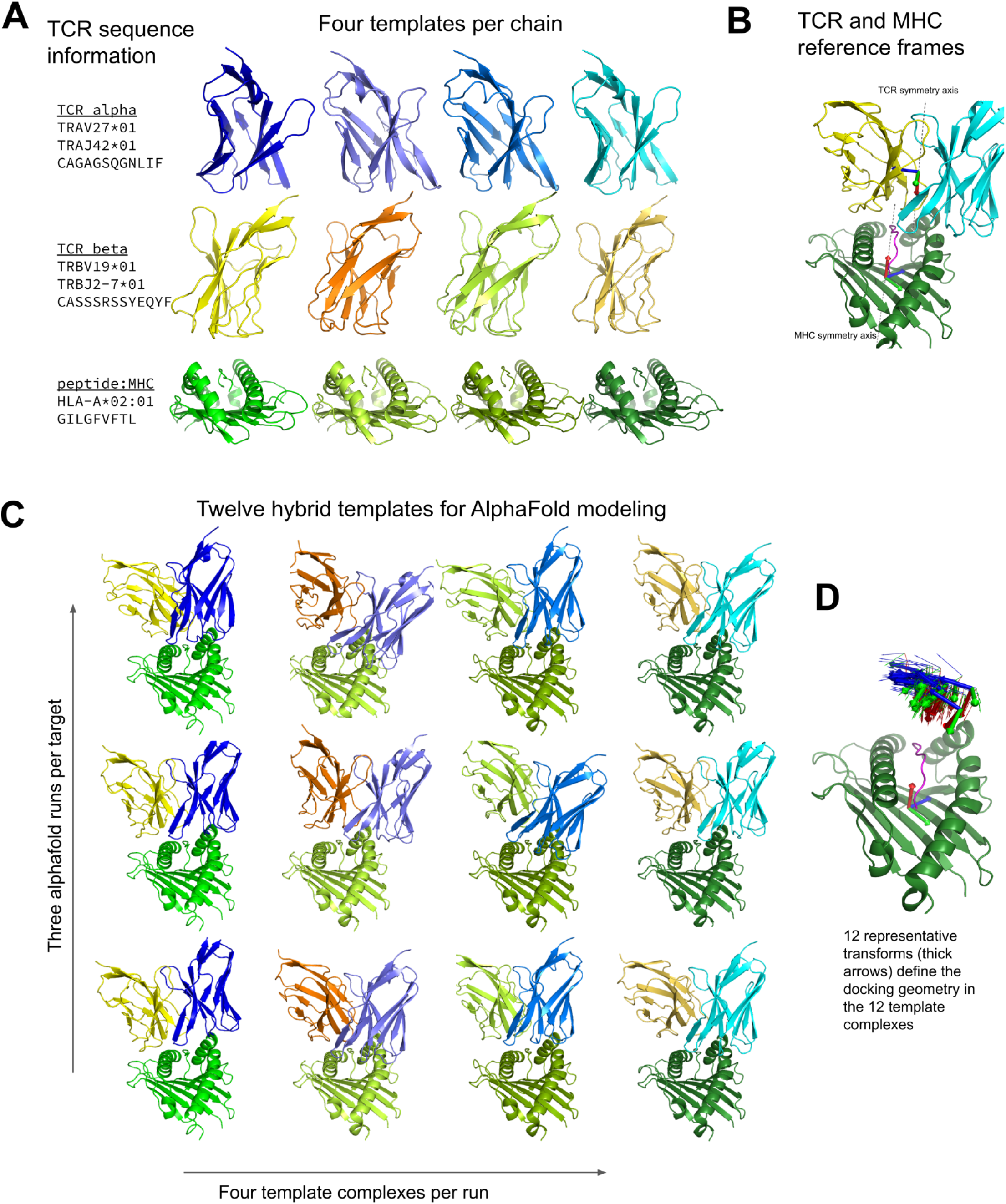
Constructing diverse hybrid templates for AlphaFold modeling. **(A)** Four structural templates for each TCR chain and for the peptide:MHC are identified in the Protein Databank (Berman et al., 2000) by sequence similarity search. **(B)** TCR:pMHC docking geometry is defined by computing the rigid-body transformation between TCR and pMHC coordinate frames. Coordinate frames are oriented based on internal pseudosymmetry as described in the Methods. **(C)** Three independent AlphaFold simulations are performed, each with four hybrid templates built from the four sets of single-chain templates oriented relative to one another using one of twelve representative docking geometries chosen to cover a wide range of experimentally determined ternary complexes. **(D)** TCR coordinate frames from class I pMHC ternary structures and the 12 representative transforms (thicker arrows) are shown in a common coordinate system defined by their corresponding pMHC coordinate frames.

We found that the hybrid templates AlphaFold pipeline specialized for TCR:pMHC (‘AF_TCR’) produces higher quality models than either of the Alphafold-Multimer variants on a benchmark set of 130 TCR:pMHC complexes (**Fig. 2A,** Wilcoxon *P*<10^-7^ vs AFM_full and *P*<10^-12^ vs AFM_trim on the full set, and **Fig. 2B**, *P*<10^-3^ for both comparisons on 20 targets without a close homolog in the AlphaFold-Multimer training set). There was a strong positive correlation between predicted and observed model accuracy (**Fig. 2C**). An attractive feature of neural network architectures is the potential to ‘fine tune’ a general network for improved prediction accuracy in a specific domain. We fine-tuned the AlphaFold parameters in the context of the AlphaFold TCR pipeline on the set of 93 human TCR:pMHC complexes from the benchmarking set and subsequently evaluated the performance of this model on the 37 mouse TCR:pMHC targets. Despite the small size of the TCR:pMHC ternary structure database, the fine-tuned model showed improved performance on the mouse targets (**Fig. 2D**; Wilcoxon *P*<0.015), which are distinct in the details of their epitope, MHC, and TCR sequences from the human training set, suggesting that the model was able to learn generalizable features of TCR:pMHC interactions. This fine-tuning procedure was facilitated by the fact that the AF2 model requires significantly less memory in the absence of MSA information, making it possible to perform parameter optimization on full TCR:pMHC systems without any residue cropping.

**Figure 2.**
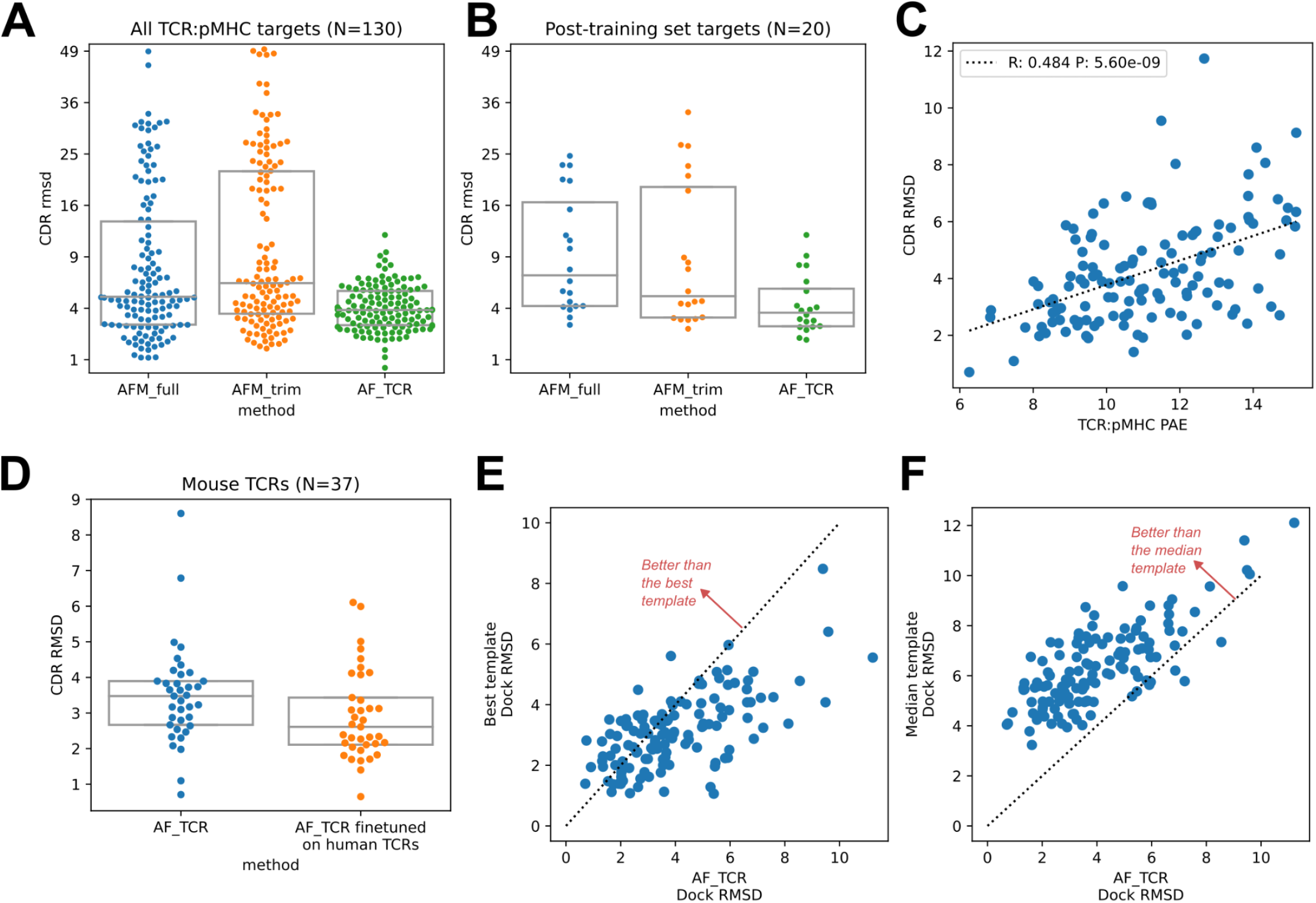
TCR modeling accuracy. **(A)** Comparison between Alphafold-Multimer with full (‘AFM_full’) or trimmed (‘AFM_trim’) input sequences and the hybrid-templates TCR pipeline (‘AF_TCR’). CDR RMSD values (y-axis) are computed by superimposing the native and modeled MHC coordinates and comparing the placement of the TCR CDR loops (see Methods). **(B)** Same as in (A) but for the 20 benchmark targets unrelated to any TCR:pMHC structure deposited before May 2018, the cutoff date for the AlphaFold-Multimer training set. **(C)** AlphaFold’s predicted aligned error (PAE) measure, evaluated between TCR and pMHC, correlates with CDR RMSD between model and native structure. **(D)** Fine-tuning AlphaFold’s parameters on human TCR:pMHC complexes improves prediction of mouse TCR:pMHC complexes. **(E)** The docking geometry of the final AlphaFold model improves over the best of the 12 templates in 30% of cases (points above the line *y*=*x*). **(F)** The docking geometry of the final AlphaFold model improves over the median of the 12 templates in 94% of cases (points above the line *y*=*x*).

For each benchmark target, the AlphaFold TCR pipeline is provided with 12 hybrid template complexes whose TCR:pMHC docking modes are taken from 12 diverse ternary structures unrelated to the target. We were curious to know whether the AlphaFold simulation was improving on the docking information present in these template structures. To answer this question, we compared the accuracy of the docking geometry present in the final model to the accuracies of the 12 template structures. Since the 12 templates differ in the sequences and structures of their CDR loops, we developed a distance between TCR:pMHC docking geometries that compares the placement of ‘generic’ CDR loops (‘docking RMSD’, see Methods). This docking RMSD measure is correlated with CDR RMSD in comparisons of models to natives (**Fig. S2**), but it focuses exclusively on the docking geometry and provides a sequence-independent way of comparing binding modes that emphasizes CDR loop placement. For 30% of the targets, the AlphaFold TCR final model had a lower RMSD than the best template docking geometry (**Fig. 2E**); the final model improved over the median template RMSD for 94% of the targets (**Fig. 2F**). To visualize the overall docking geometry landscape of models and natives, we calculated docking RMSD values between all of the native ternary structures and the AlphaFold-TCR and AlphaFold-Multimer models and transformed this distance matrix into a 2D projection (**Fig. S1**) using the UMAP algorithm (McInnes et al., 2018). Inspection of this 2D docking geometry landscape reveals regions that are distant from the native structures and only sampled by the AlphaFold-Multimer models, supporting the view that incorporating template docking geometries helps to constrain predictions to native-like geometries.

### Binding specificity prediction

Having established that the AlphaFold TCR pipeline can generate more accurate TCR:pMHC models than AlphaFold-Multimer, we evaluated its performance in TCR epitope prediction. The general problem of predicting, de novo, which peptide:MHCs a given TCR recognizes is likely to be very difficult due to the diversity of TCR:pMHC recognition modes, the polyspecificity of individual TCRs, and the paucity of available training data (Moris et al., 2021). Here we consider instead the simpler problem of selecting the correct target peptide from a small set of candidates. This might correspond to a real-world scenario in which we know the source antigen from which the unknown peptide epitope is taken, or we have a positive hit in a T cell stimulation assay that implicates a pool of peptides rather than a unique epitope. For benchmarking, we focus on peptide-MHC epitopes for which a repertoire of cognate TCRs has been identified. This allows us to evaluate the sensitivity of the predictions to small changes in TCR sequence. It also lets us investigate a scenario in which we are given not one TCR, but a set of TCRs that all recognize the same epitope, and we consider the extent to which this helps to constrain the target epitope. With improved single-cell technologies for paired TCR sequencing, and improved methods for identifying TCR sequence convergence, we hypothesize that this will become an increasingly common scenario.

We selected a set of 8 Class I peptide:MHC systems (**Table 1**) for which a minimal repertoire of paired epitope-specific TCRs and a solved ternary structure were available. These systems include one human (A*0201) and one mouse (H2-Db) MHC allele, each with 9- and 10-residue peptides. TCR repertoires containing more than 50 unique TCR sequences were subsampled to a set of 50 TCRs using an algorithm that removed redundancy while concentrating on the more densely sampled regions of TCR space (see Methods). For each MHC/peptide length combination, we used the NetMHCpan-4.1 (Reynisson et al., 2020) method to select 9 decoy peptides with binding scores in the range of the true peptide binders. We additionally selected 50 irrelevant TCRs at random from human and mouse CD8 T cell datasets made available by 10X Genomics (see Methods).

**Table 1.** Binding specificity benchmark

| Organism | MHC | Peptide length | Peptide sequence | Antigen |
| --- | --- | --- | --- | --- |
| human | HLA-A*02:01 | 9 | GILGFVFTL | Flu M1 |
| human | HLA-A*02:01 | 9 | GLCTLVAML | EBV BMLF1 |
| human | HLA-A*02:01 | 9 | NLVPMVATV | CMV pp65 |
| human | HLA-A*02:01 | 9 | YLQPRTFLL | SARS-CoV-2 Spike |
| human | HLA-A*02:01 | 10 | ELAGIGILTV | human MART-1 |
| human | HLA-A*02:01 | 10 | KLVALGINAV | HCV POLG |
| mouse | H2-Db | 9 | ASNENMETM | Flu NP |
| mouse | H2-Db | 10 | SSLENFRAYV | Flu PA |

We used the AlphaFold TCR pipeline to generate docked complexes and associated interface accuracy estimates for pairings of each TCR with its true pMHC epitope and with 9 decoy peptides of the same length (**Fig. 3A**). This produces, for each of the eight pMHCs, an Nx10 matrix of predicted interface accuracies (**Fig. 3B**, left panel), where N is the number of TCRs specific for the given pMHC. To generate a single number representing the estimated interface accuracy of a complex, we summed the residue-residue predicted aligned error (PAE) for all TCR:pMHC residue pairs. These raw accuracy estimates showed significant TCR- and pMHC-intrinsic effects (**Fig. 3B**). Certain TCRs had consistently higher or lower than average predicted interface accuracies due to features such as longer CDR3 loops or usage of V genes without a close structural template. We saw similar, albeit weaker, trends across different peptide:MHC complexes, perhaps due to AlphaFold’s confidence in the MHC-bound structure of the peptide. TCR-intrinsic factors do not change the relative order of candidate peptides, but they make comparisons across TCRs difficult; pMHC effects have the potential to change the rank ordering of candidate peptide epitopes. Since we are interested here in evaluating the compatibility between TCR and pMHC and not, for example, ranking peptides by their affinity for MHC, we corrected for these TCR- and pMHC-intrinsic effects to generate an array of TCR:pMHC binding scores intended to be comparable across different pMHCs and TCRs (**Fig. 3B**, middle panel; lower scores indicate stronger predicted binding, see Methods).

**Figure 3.**
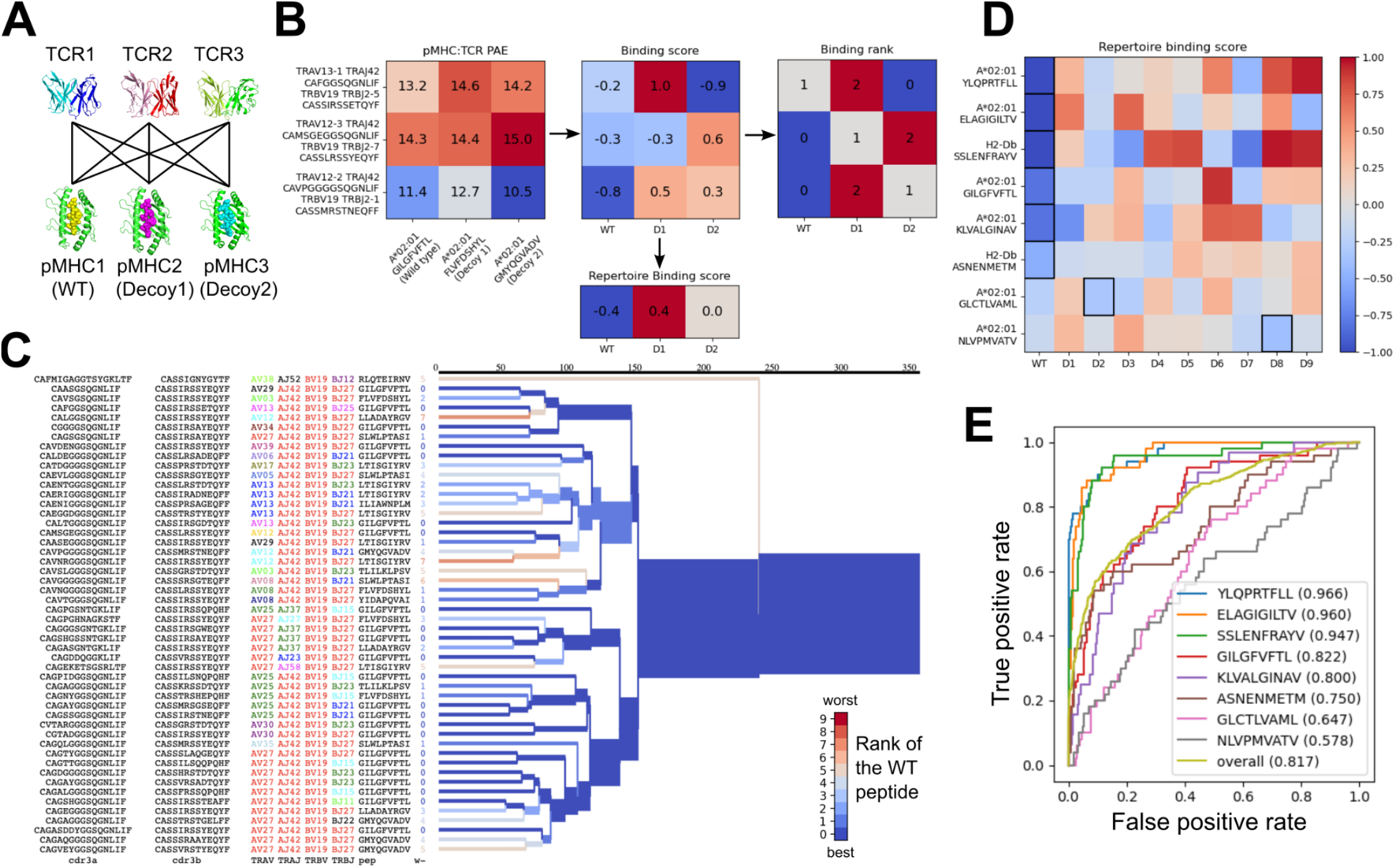
Structural modeling can discriminate correct from incorrect TCR:pMHC pairings. **(A)** For each of the eight peptide:MHC epitopes, we docked multiple cognate TCRs against multiple decoy peptides and the wild type epitope. Here three TCRs and three pMHCs are shown; 9 decoys and up to 50 TCRs were actually modeled. **(B)** For each candidate TCR:pMHC pairing, the mean AlphaFold predicted aligned error (PAE) for the TCR:pMHC interface was calculated (left) and transformed into a binding score by subtracting out TCR-intrinsic and pMHC-intrinsic factors (middle). These binding scores were averaged to define a repertoire-level binding score for the WT epitope and each of the decoys (bottom). Also calculated was the rank of the WT binding score within the list of all the binding scores for each TCR (right). **(C)** TCRdist hierarchical clustering tree of the 50 modeled TCRs for the A*02:01 GIL9 epitope, labeled with the TCR sequence information, top-ranked peptide, and rank of the WT peptide, and colored by the rank of the WT peptide. Internal edges, which correspond to multiple “leaf” TCRs, are colored by the rank of the WT peptide after averaging the binding scores over the leaf TCRs. **(D)** Repertoire binding scores for each of the eight target epitopes and the 9 decoy peptides, with the lowest (most favorable) binding score in each row boxed. **(E)** Receiver operating characteristic (ROC) curves for discrimination of WT from decoy peptides by binding score. Area under the ROC curve (AUROC) values are given in the legend along with the sequence of the WT peptide.

We evaluated the accuracy of these binding predictions across the eight pMHC epitopes. First, we calculated the rank of the true peptide epitope amongst the 9 decoy peptides (**Fig. 3B**, right panel) on a per-TCR basis. To visualize how these ranks vary across each pMHC-specific repertoire, we constructed hierarchical clustering trees of the TCR sequences using the TCRdist measure (Dash et al., 2017) and colored them by the rank of the true peptide (**Fig. 3C** and **Fig. 4**). Internal edges, which correspond to multiple “leaf” TCRs, are colored by the rank of the true peptide after averaging the binding scores over the leaf TCRs. Looking across all eight epitopes, we can see, first, that the predictions are not random: on average the correct peptide is ranked more favorably than most of the decoys (i.e., there is more blue than red). For six of the eight epitopes, the correct peptide is ranked first when we average the binding scores of all the TCRs in the repertoire (**Fig. 3D**; **Fig. 4**: the largest branch of the tree is dark blue). It also appears that the epitopes with more sequence-diverse repertoires (A*0201-GLC9 and A*02:01-NLV9) are more challenging to predict: the trees that merge completely at smaller TCRdist values (further to the left) are bluer than the other trees in **Figure 4**. This can be seen quantitatively by plotting the TCRdiv repertoire sequence diversity measure (Dash et al.,2017) against measures of binding prediction success (**Fig. S3**; **Algorithm S1**). If we rank the peptides by binding score and compare the recovery of true binder peptides to decoys using receiver operating characteristic (ROC) curves, we can see that some epitopes, such as A*02:01-YLQ9 and A*02:01-ELA10 are predicted very well (by area under the ROC curve, AUROC >= 0.96) and some predictions are only slightly better than random (**Fig. 3E**). We find an overall AUROC value of 0.82 when binding and non-binding TCR:pMHC pairs from all epitopes are ranked together.

**Figure 4.**
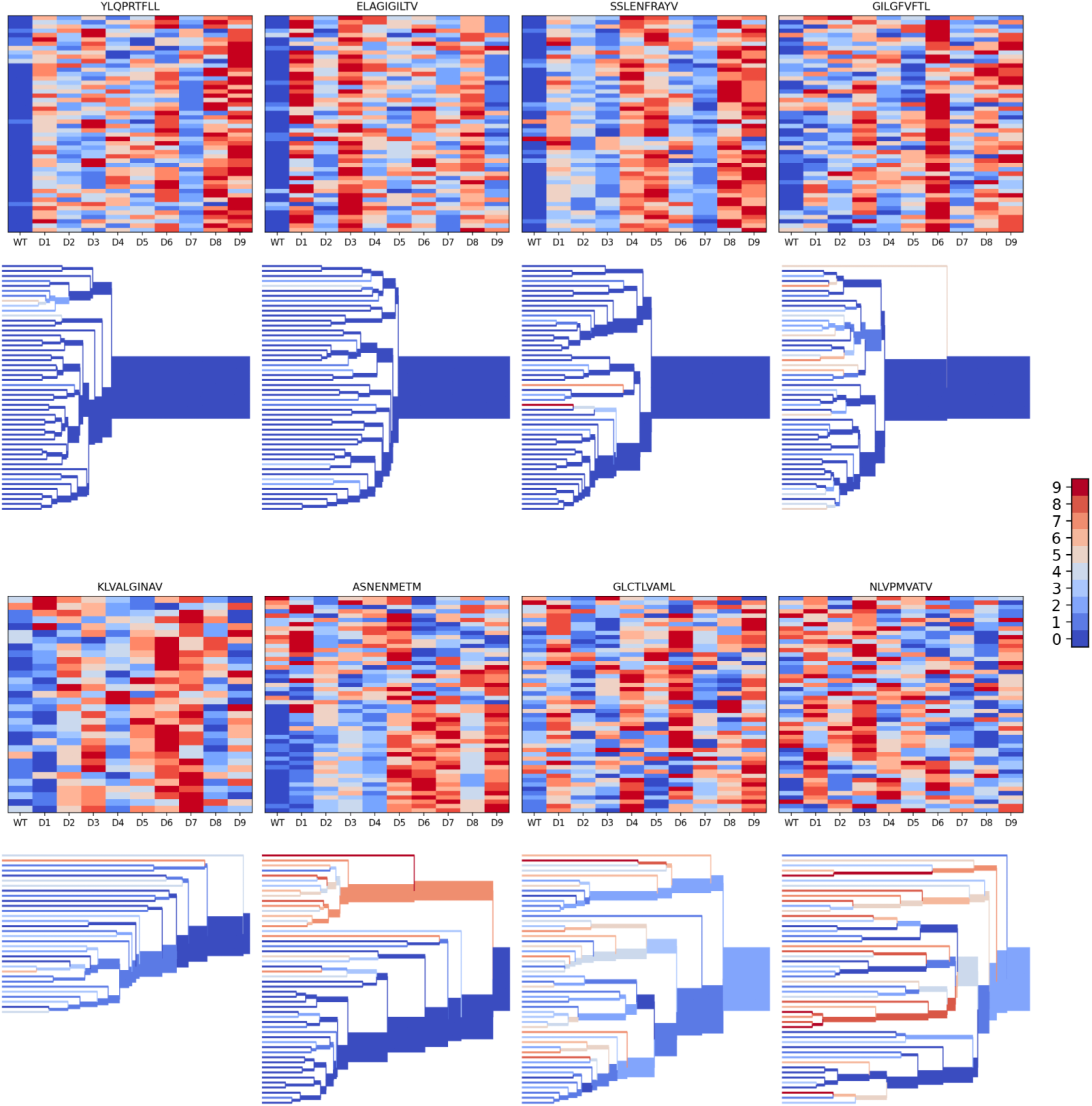
Peptide decoy discrimination results for the eight benchmark epitopes. The rank of the wild type peptide relative to the 9 decoys (0=best, 9=worst) is shown in a heatmap and a TCRdist hierarchical clustering tree of the epitope-specific TCRs. Each row of the heatmap corresponds to a single TCR; each column corresponds to one of the 10 modeled peptides, with the wild type peptide on the left. The vertical ordering of the TCRs in the heatmaps and trees is the same. Internal edges of the trees, which correspond to multiple “leaf’ TCRs, are colored by the rank of the wild type peptide after averaging the binding scores over the leaf TCRs.

We looked to see whether structural modeling accuracy correlated with binding prediction success (**Fig. 5**). Although very few of the specific TCRs being modeled have been structurally characterized, each of the epitopes has at least one solved ternary structure in the protein structure database. For each TCR, we computed docking RMSDs between the TCR:pMHC model in complex with its cognate epitope and the solved ternary structures for that epitope and took the minimum value as a proxy for the accuracy of the predicted binding mode. **Figure 5A** shows the distribution of these RMSD values across each repertoire. Well-predicted epitopes such as A*02:01-YLQ9 and A*02:01-ELA10 indeed appear to have smaller RMSD values than other repertoires. The mouse pMHC H2Db-ASN9 is an outlier, with an RMSD distribution shifted to very high values. Examination of the three ternary structures for this pMHC revealed that they represent a unique population of TRBV17+ TCRs that is distinct from the consensus repertoire modeled here. Two of the three TCRs bind with a reversed docking orientation, and the third has a highly displaced binding footprint (Zareie et al., 2021); all three are outliers in a hierarchical clustering tree of Class I TCRs based on docking RMSD (**Fig. S4**). If we exclude H2Db:ASN9 and plot docking RMSD to the closest epitope structure versus binding score for the correct peptide, we see that there is a positive correlation (**Fig. 5B**). The TCRs for which the correct peptide is ranked first have a lower RMSD distribution than other TCRs, and this RMSD distribution shifts upward as the rank of the correct peptide declines (**Fig. 5C**). These results suggest that the correct binding predictions are driven at least in part by recovery of native-like structural features.

**Figure 5.**
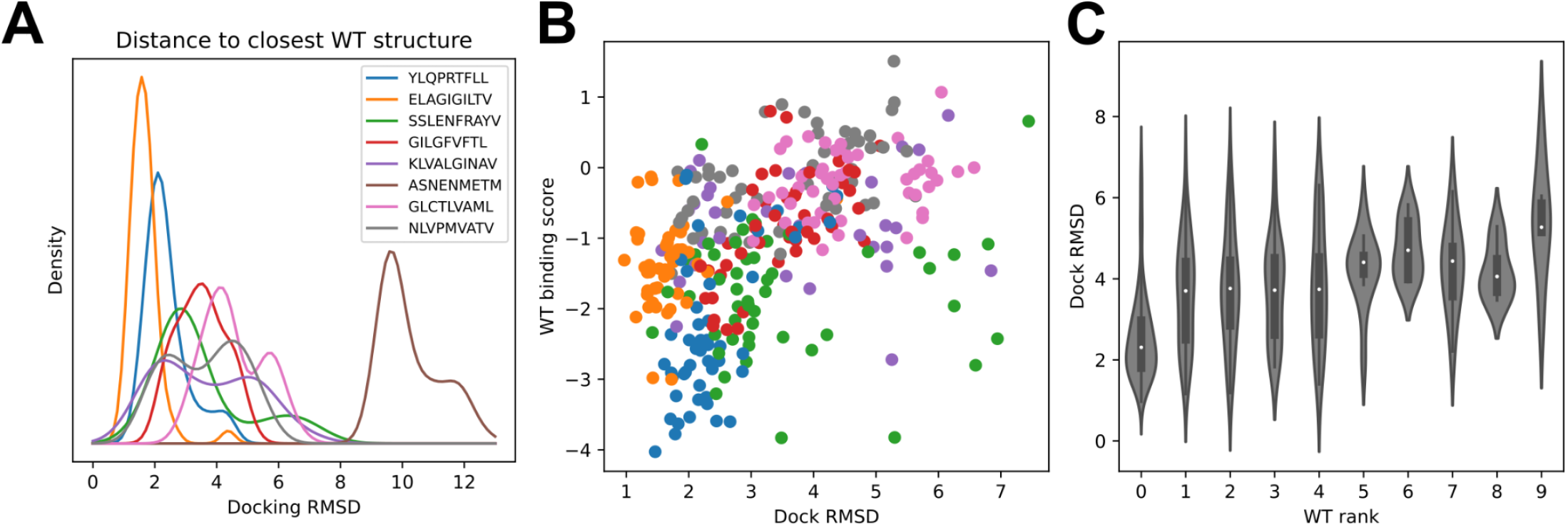
Success in decoy discrimination correlates with structural modeling accuracy. **(A)** For each TCR, the structural model in complex with the wild type epitope was compared to all experimentally determined ternary structures for that epitope and the smallest docking RMSD was recorded. The resulting RMSD distributions were smoothed using kernel density estimation and plotted. **(B)** Scatter plot of docking RMSD to the nearest wild type structure versus the binding score for the wild type peptide. Favorable wild type binding scores correlate with lower RMSD values. **(C)** Distributions of docking RMSD to the nearest wild type structure (y-axis) as a function of the rank of the wild type peptide (x-axis). When the wild type peptide is ranked first (left violin), the corresponding docking geometries are more similar to those of ternary complexes for that epitope, suggesting higher accuracy.

To further investigate the specificity of our modeling approach, we performed an *in silico* epitope alanine scan of each of the eight pMHC-specific repertoires. We built models and calculated binding scores for each epitope-specific TCR docked to all single-alanine mutants of the native peptide (native alanine residues were mutated to glycine). Binding scores for each TCR and each of the alanine mutants are shown in the heatmaps in **Figure 6**. Averaging these binding scores over all TCRs for each epitope and subtracting the score for the native peptide gives a predicted repertoire-level sensitivity to mutation at each peptide position (**Fig. 6B**). From these sensitivity plots we can see that the majority of the epitope-specific repertoires show the expected preference for the native peptide at most positions, with a subset of positions showing high sensitivity. Coloring the pMHC structures by mutation sensitivity (**Fig. 6A**) reveals that these highly sensitive positions are largely TCR-exposed. The two pMHCs with poor AUROC values for decoy discrimination (A*02:01-GLC9 and A*02:01-NLV9) also show weak specificity for the native peptide here.

**Figure 6.**
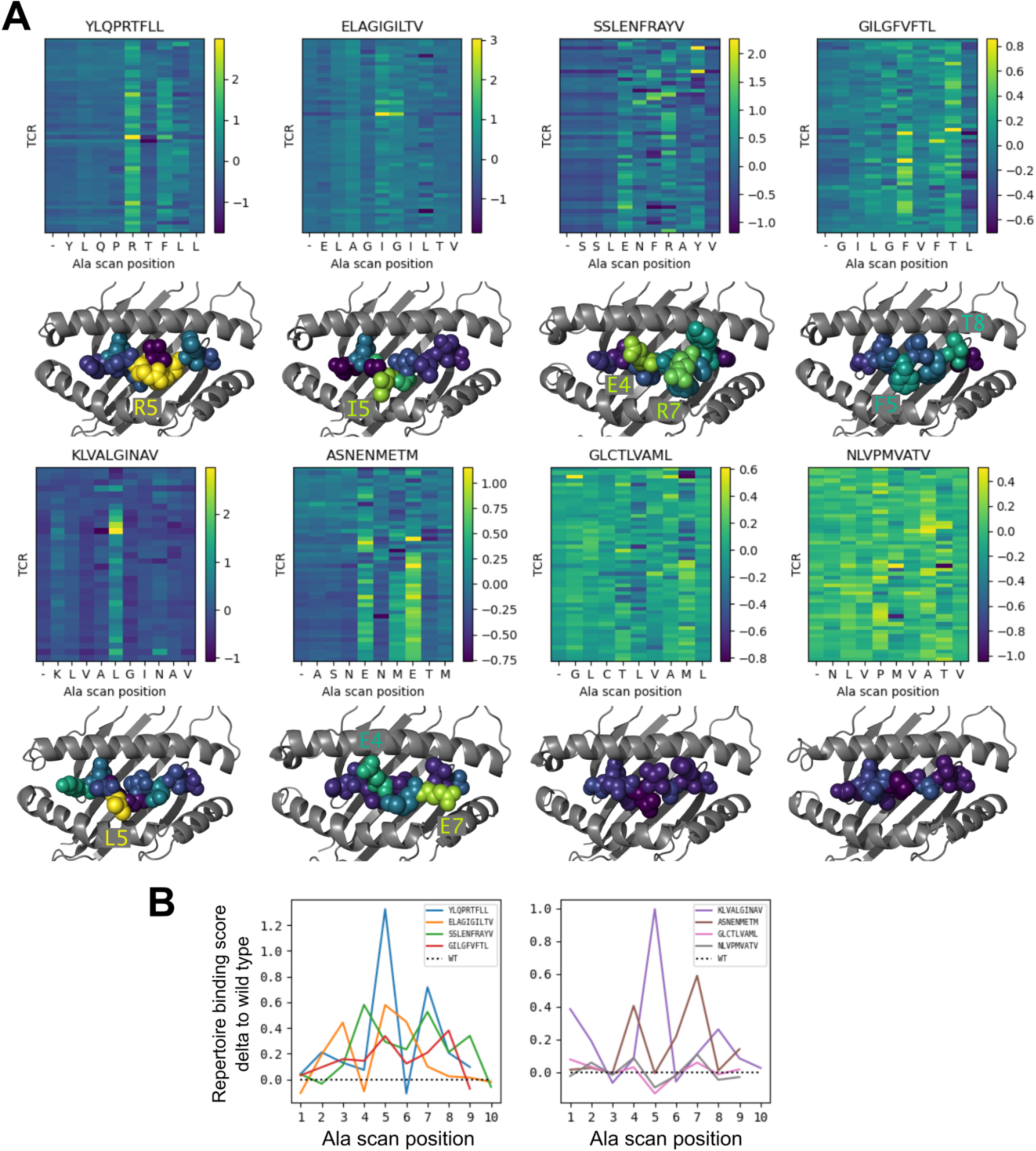
Alanine scanning results for the eight benchmark epitopes. **(A)** Heatmaps showing the binding scores for the wild type peptide (left column) and all single-alanine mutants (columns labeled with the wild type sequence) in complex with each TCR (rows). Below each heatmap, the wild type pMHC crystal structure is shown with the peptide colored by the delta between mutant and wild-type repertoire-averaged binding scores. **(B)** Line plots of the delta between the mutant and wild-type repertoire-averaged binding scores reflect the predicted repertoire-level sensitivity to epitope mutations.

## Discussion

Prediction of TCR:pMHC interactions is challenging because of the diversity of TCR:pMHC recognition modes and the limited number of validated interactions available for training. Inspired by recent breakthroughs in protein structure prediction (Baek et al., 2021; Jumper et al., 2021), we hypothesized that structure-based approaches, which can leverage general features of protein structures and interactions, might offer a path to generalizable TCR:pMHC binding predictions from limited data. We developed a specialized AlphaFold pipeline for TCR:pMHC structure prediction that uses hybrid templates assembled from existing TCR:pMHC structures to constrain the TCR docking orientation to native-like geometries. Here we show that this pipeline can generate more accurate structure predictions of TCR:pMHC complexes than the state-of-the-art method Alphafold-Multimer. Prediction accuracy correlates with model confidence, and model quality can be further improved by fine-tuning the AlphaFold parameters on TCR:pMHC structures. When tested on peptide decoy discrimination, we found that the model’s docking accuracy estimates, corrected for TCR- and pMHC-intrinsic effects, could be used to select the correct target peptides from decoys with substantial accuracy. Success in this decoy discrimination task correlated with the structural accuracy of the models, suggesting that the pipeline was picking out the correct peptide on the basis of molecular specificity determinants. Prediction accuracy varied across pMHC epitopes, with those epitopes having more sequence-diverse TCR repertoires proving more challenging to model.

There are a number of caveats to this work. First, the overall level of accuracy falls short of what would be required for most practical applications of TCR:pMHC binding prediction. As described below, we are pursuing multiple avenues for improving this initial pipeline; it may also be possible to predict from the simulations themselves which systems are reliably modeled, which could allow useful predictions to be extracted from large-scale calculations. Second, several of the epitopes in our peptide decoy discrimination benchmark have been extensively characterized in structural studies. While we made efforts to avoid using information from related structures during template assembly (see Methods), it is still possible that bias toward native-like conformations was introduced. For example, the AlphaFold parameters we rely on in the pipeline were trained on individual protein chains (not protein complexes) deposited prior to May, 2018. Some of the TCR chains modeled in the decoy discrimination task are likely similar to protein chains present in this AlphaFold training set. Finally, our template-based modeling approach is unlikely to succeed on TCR:pMHC systems with highly divergent binding modes. Although we do see evidence that AlphaFold can improve over the best template provided (**Fig. 2E**), it is unlikely that it can reliably predict complexes that deviate substantially from any template (for example, reversed-orientation geometries (Zareie et al., 2021)).

The modeling pipeline described here represents a first step in applying deep learning structure prediction tools to study TCR:pMHC interactions. We anticipate that it can be improved by further testing on other systems and by comparison with other experimental data types (binding affinities, interface mutations, etc). This initial pipeline does not make use of multiple sequence alignment (MSA) information, but it may be helpful to include MSAs for individual chains or to construct “paired MSAs” consisting of concatenated TCR:peptide:MHC sequences of known binding examples. Such paired MSAs could take the place of the paired ortholog alignments used by AlphaFold-Multimer to detect residue covariation across interfaces. We evaluated the use of AlphaFold’s residue-residue accuracy estimate (PAE) to discriminate wild type from decoy peptide-MHC epitopes, but it may also be worth exploring the use of other binding affinity estimates such as binding energies computed with the

Rosetta software package (Leaver-Fay et al., 2011) or other molecular modeling tools (Lee et al.,2018). Finally, it may be possible to fine-tune AlphaFold parameters directly to discriminate TCR:pMHC binding examples from non-binding examples, as we have recently demonstrated for peptide:MHC interactions (Motmaen et al., 2022). This would allow us to directly leverage the thousands of validated TCR:pMHC interactions within the context of a structurally-informed training procedure.

## Methods

### Defining TCR:pMHC docking geometry

The TCR:pMHC docking geometry is defined by the rigid body transformation that maps between the MHC and TCR coordinate frames (**Fig. 1B**). The MHC coordinate frame is defined on the basis of the approximate 2-fold symmetry axis that relates the N- and C-terminal halves of the beta sheet forming the floor of the peptide binding pocket. 12 core residues in the beta sheet were selected (**Fig. S5A**), 6 from the N-terminal half and 6 from the C-terminal half, that are related by this approximate 2-fold rotational symmetry. For a given MHC structure, the transformation mapping these 12 residues onto themselves, interchanging the N- and C-terminal residues and minimizing the RMSD of the alpha carbon atoms, is computed. The rotation axis of this orthogonal transformation, oriented to point toward the peptide, is taken as the x-axis of the MHC coordinate frame. The z-axis of the coordinate frame points from the center of mass (COM) of the 6 N-terminal core alpha carbons to the COM of the 6 C-terminal core alpha carbons. The coordinate frame is centered at the COM of the 12 core residues.

To define the TCR coordinate frame, 13 structurally conserved core residues from the TCR alpha chain and 13 aligned core residues from the TCR beta chain (**Fig. S5B-C**) were selected on the basis of visual inspection of TCR multiple structural alignments. The same procedure as outlined above for the MHC is used to define the TCR coordinate frame, replacing the 6 N-terminal and 6 C-terminal core residues of the MHC with the 13 TCRA and 13 TCRB core residues of the TCR heterodimer. The Python script `parse_tcr_pmhc_pdbfile.py` in the TCRdock github repository (see Code Availability) computes the MHC and TCR coordinate frames for an input PDB structure and calculates the docking geometry.

### AlphaFold modeling pipeline

To model a given TCR:pMHC target, three AlphaFold simulations (using the `‘ model_2_ptm’` parameter set) are conducted and the final model with the lowest predicted aligned error (PAE) between the TCR and pMHC is selected (**Figure 1**). To reduce parameter training bias, we used the original AlphaFold monomer parameters, which were trained on single protein chains, rather than the AlphaFold-Multimer parameter set, whose training set included protein complexes. Each AlphaFold simulation can use a maximum of four templates, allowing for 12 total templates across the three runs (**Fig. 1C**). These 12 templates are constructed from four templates for each of the pMHC, TCRA, and TCRB chains selected on the basis of sequence identity to the modeling target (**Fig. 1A**). Peptide-MHC templates are sorted by total sequence identity computed over both the MHC and the peptide. To create hybrid templates for AlphaFold modeling, the pMHC and TCRB template coordinates must be mapped into the coordinate frame of the TCRA template structure. First, the TCR structure from which the TCRB template coordinates are being taken is superimposed onto the TCRA template structure by superimposing the 13 TCRA core residues. Then the superimposed TCRB coordinates are appended to the hybrid template after the TCRA coordinates. To map the pMHC coordinates into the coordinate frame of the TCRA+TCRB coordinates, MHC and TCR coordinate frames are defined as described above and 12 representative docking geometries are selected. Each docking geometry defines the transformation between the MHC and TCR coordinate frames, allowing the pMHC template coordinates to be mapped into the hybrid template TCR coordinate frame. To choose the 12 representative docking geometries, docking geometries from TCR:pMHC structures of the same MHC class as the target are hierarchically clustered and the clustering tree is cut at a distance threshold at which there are 12 clusters. The docking geometry from each cluster with the smallest mean distance to the other cluster members is chosen as the representative. For hierarchical clustering, a matrix of docking RMSDs (defined below) is provided to the `hierarchy. linkage` function in the SciPy (Virtanen et al., 2020) `cluster` module. The `hierarchy. fcluster` function with ‘maxclust’ criterion is used to select the distance threshold at which the docking geometry tree divides into 12 clusters. Template structures were downloaded from the RCSB Protein Databank (Berman et al., 2000) ftp site on 2021-08-05.

### Fine-tuning AlphaFold for TCR:pMHC structure prediction

To fine tune the AlphaFold parameters for TCR:pMHC structure prediction, we used a version of the AlphaFold package that was modified slightly to expose the parameter training interface (Motmaen et al., 2022). The Python script `run_finetuning_for_structure.py` in the `alphafold_finetune` github repository (https://github.com/phbradley/alphafold_finetune) with the additional command line flags `‘--model_name model_2_ptm --crop_size 419’` was provided with a training set consisting of three runs for each of the 93 human ternary structures (279 total training examples). Due to the small size of the training dataset, training was stopped after two epochs to avoid over-fitting.

### Structure prediction benchmark

The structure prediction benchmark set consists of 130 nonredundant ternary TCR:pMHC structures deposited prior to 2021-08-05 (**Table S1**). No two structures in the set have fewer than 3 peptide mismatches *and* a paired TCRdist (Dash et al., 2017) distance less than or equal to 120. This constraint eliminates pairs of structures with the same or similar TCRs binding to the same or similar peptides. After visual inspection, we eliminated the following 9 outlier structures with highly divergent binding modes (reversed docking orientations, extremely bulged peptides, etc.): PDB IDs 5sws, 7jwi, 4jry, 4nhu, 3tjh, 4y19, 4y1a, 1ymm, and 2wbj.

During benchmarking, we excluded templates and docking geometries that were too similar to the target sequence being modeled. Peptide-MHC templates were excluded if they had fewer than 3 peptide mismatches with the target peptide. TCR chain templates were excluded if they had a single-chain TCRdist of 36 or less to the target chain (corresponding to 3 non-conservative mismatches or indels in the CDR3 loop). Docking geometries were excluded if they came from a structure with fewer than 3 peptide mismatches to the target or a TCRdist of 48 or less from the target TCR.

### RMSD measures

We assessed model accuracy by comparing the placement of the CDR loops relative to the MHC in the native and modeled structures. The two structures were first superimposed on the MHC coordinates; then an alpha-carbon RMSD was calculated (without further superposition) over the CDR loops, up-weighting residues in the CDR3 by a factor of 3 to reflect the greater importance of the CDR3 for epitope recognition (this is the ‘CDR RMSD’ reported in **Figure 2**). TCRdist CDR loop definitions were used.

To compare docking geometries between structures with different CDR loop sequences, we developed a ‘docking geometry RMSD’ intended to approximate the CDR RMSD in a sequence-independent fashion. The full template database was first used to calculate a mean center of mass of the residues in each CDR loop with respect to the TCR coordinate frame. To compute the docking RMSD between two docking geometries, each docking geometry is used to build a TCR coordinate frame assuming the MHC coordinate frame is centered at the origin and aligned with the coordinate axes. Then the CDR centers of mass are built with respect to each of these two TCR coordinate frames, and an RMSD is calculated between these two sets of eight points (4 CDR centers of mass each for the TCRA and TCRB chains) without superposition, upweighting the CDR3 center of mass by a factor of 3. The correlation between CDR RMSD and docking RMSD is shown in **Figure S2**.

### Decoy discrimination benchmark

Eight MHC class I epitopes with TCR repertoire data and experimentally determined structures were selected as targets for a decoy discrimination benchmark (**Table 1**). Paired alpha and beta sequences of TCRs specific for these eight epitopes were collected from the literature (10x_Genomics, 2020; Dash et al., 2017; Francis et al., 2022; Minervina et al., 2022; Schattgen et al., 2021; Shugay et al., 2018). Epitope-specific TCR repertoires with more than 50 TCRs were subsampled to 50 representatives using a Gaussian kernel density-based algorithm designed to preferentially sample denser regions of TCR space without introducing excessive redundancy (**Algorithm S2**). The goal in sampling denser regions of TCR space was to avoid outlier TCR sequences that might represent experimental errors. 100 additional ‘irrelevant’ TCR sequences (50 mouse TCRs and 50 human TCRs) were selected at random from naive CD8 T cells in datasets made publicly available by 10X Genomics (https://www.10xgenomics.com/resources/datasets). All TCR sequences are listed in **Table S2**.

The eight MHC class I epitopes include 9 and 10 residue peptides presented by the MHC alleles HLA-A*02:01 and H2-Db. For each MHC and peptide length, 9 decoy peptides were selected by scanning a 1500 residue artificial source antigen sequence with NetMHCpan-4.1 (Reynisson et al.,2020) and selecting the top 9 predicted binders. The artificial source antigen sequence was created by concatenating the source antigen sequences for the 9 benchmark targets (**Table 1**), shuffling, and selecting the first 1500 residues.

Each epitope-specific TCR was modeled in complex with its cognate peptide epitope and in complex with the 9 length- and MHC-matched decoy peptides using the AlphaFold pipeline specialized for TCRs. The mean predicted aligned error (PAE) residue-residue accuracy measure for TCR:pMHC residue pairs was calculated for each complex and stored in an Nx10 matrix, where N is the number of TCRs (each row corresponds to a TCR and each column to a peptide). To convert these raw TCR:pMHC PAE values into a binding score that can be compared across TCRs and pMHCs, we also modeled each pMHC in complex with 50 irrelevant background TCRs from the same organism. The mean TCR:pMHC PAE for these background complexes was calculated for each pMHC and was subtracted from the matrix column of PAE values involving that pMHC. The values in the resulting matrix of adjusted PAE values were then shifted to have 0 row sums by subtracting its mean value from each row. Thus in the final Nx10 matrix of binding scores, the mean value for each row is 0, while the mean values of the columns reflect the overall binding preference of the full repertoire of TCRs for the peptide corresponding to the column.

During modeling, the TCR- and pMHC-similarity constraints described above in ‘Structure Prediction Benchmark’ were applied to exclude templates; in addition, ternary structures with a peptide having fewer than 3 mismatches from the wildtype peptide were excluded from all simulations (with decoy or wildtype peptides).

The epitope alanine-scanning benchmark was performed as described above with the difference that the decoys were single-residue alanine mutants of the wild type peptide (alanine residues in the wild type peptide were mutated to glycine). Thus there were 9 decoys for 9-residue peptides and 10 decoys for 10-residue peptides.

## Software and Data Availability

Python software to set up and run the TCR-specialized AlphaFold pipeline described here and to parse TCR:pMHC ternary structures is available in the TCRdock github repository (https://github.com/phbradley/TCRdock). Benchmark datasets are provided as Supplementary Data attached to this manuscript and are also available in the TCRdock github repository.

## Acknowledgments

I am grateful to Jeremy Crawford, Anastasia Minervina, Amir Motmaen, Paul Thomas, and Albert Yeh for helpful comments on the manuscript, to Justas Dauparas for help fine-tuning AlphaFold, to the creators of AlphaFold for freely sharing their software and parameters, and to Fred Hutch Scientific Computing and NIH ORIP S10OD028685 for outstanding computing infrastructure. This research was supported by NIH grants R35 GM141457 and R01 AI136514.

