## Supplementary Data for "Structure-based prediction of T cell receptor:peptide-MHC interactions"

### Supplementary Material

##### Supplementary Table 1. Structure prediction benchmark

| pdbid | organism | mhc_class | mhc | peptide | va | ja | cdr3a | vb | jb | cdr3b | cdr_rmsd | cdr_rmsd_af | cdr_rmsd_af2_trim |
|---|---|---|---|---|---|---|---|---|---|---|---|---|---|
| 5bs0 | human | 1 | A*01:01 | ESDPIVAQY | TRAV21*01 | TRAJ28*01 | CAVRPGGAG | TRBV5-1*01 | TRBJ2-7*01 | CASSFNMAT | 4.94 | 30.43 | 33.37 |
| 5brz | human | 1 | A*01:01 | EVDPIGHLY | TRAV21*01 | TRAJ28*01 | CAVRPGGAG | TRBV5-1*01 | TRBJ2-7*01 | CASSFNMAT | 4.45 | 29.81 | 38.26 |
| 4eup | human | 1 | A*02:01 | ALGIGILTV | TRAV12-2*02 | TRAJ45*01 | CAVSGGGAD | TRBV28*01 | TRBJ2-1*01 | CASSFLGTGV | 2.68 | 2.28 | 2.26 |
| 3utt | human | 1 | A*02:01 | ALWGPDPAAA | TRAV12-3*01 | TRAJ12*01 | CAMRGDSSY | TRBV12-4*0 | TRBJ2-4*01 | CASSLWEKLA | 6.9 | 30.25 | 19.82 |
| 5nht | human | 1 | A*02:01 | ELAGIGILTV | TRAV12-2*02 | TRAJ45*01 | CAVGGGADG | TRBV19*01 | TRBJ2-2*01 | CASSQGLAG | 2.13 | 2.66 | 3.03 |
| 3qdg | human | 1 | A*02:01 | ELAGIGILTV | TRAV12-2*01 | TRAJ23*01 | CAVNFGGGK | TRBV6-4*01 | TRBJ1-1*01 | CASSLSFGTE | 2.35 | 3.44 | 4.53 |
| 3hg1 | human | 1 | A*02:01 | ELAGIGILTV | TRAV12-2*01 | TRAJ27*01 | CAVNVAGKS | TRBV30*01 | TRBJ2-2*01 | CAWSETGLG | 4.08 | 4.47 | 5.98 |
| 5e9d | human | 1 | A*02:01 | ELAGIGILTV | TRAV12-2*02 | TRAJ24*02 | CAVTKYSWG | TRBV6-5*01 | TRBJ2-7*01 | CASRPGWM | 4.36 | 4.87 | 48.16 |
| 3qdm | human | 1 | A*02:01 | ELAGIGILTV | TRAV35*01 | TRAJ49*01 | CAGGTGNQF | TRBV10-3*0 | TRBJ1-5*01 | CAISEVGVGC | 6.67 | 5.35 | 4.04 |
| [illegible] | [illegible] | [illegible] | [illegible] | [illegible] | [illegible] | [illegible] | [illegible] | [illegible] | [illegible] | [illegible] | [illegible] | [illegible] | [illegible] |

...

https://github.com/phbradley/TCRdock/blob/main/datasets_from_the_paper/table_S1_structure_benchmark_complexes.csv

##### Supplementary Table 2. TCR sequences for the specificity prediction benchmark

| organism | mhc | wt_peptide | va | ja | cdr3a | vb | jb | cdr3b | wt_binding_score |
|---|---|---|---|---|---|---|---|---|---|
| human | A*02:01 | YLQPRTFLL | TRAV12-1*0 | TRAJ15*01 | CVVNEEDALI | TRBV20-1*0 | TRBJ2-2*01 | CSVSRDRNT | -3.236 |
| human | A*02:01 | YLQPRTFLL | TRAV12-1*0 | TRAJ30*01 | CVVNKDDKII | TRBV7-8*01 | TRBJ2-2*01 | CALSDQNTG | -3.537 |
| human | A*02:01 | YLQPRTFLL | TRAV12-1*0 | TRAJ30*01 | CVVNKEDKII | TRBV13*01 | TRBJ2-2*01 | CATLTDRNT | -3.563 |
| human | A*02:01 | YLQPRTFLL | TRAV12-1*0 | TRAJ30*01 | CVVNRDDKII | TRBV4-3*01 | TRBJ1-1*01 | CASSHRNTE | -3.592 |
| human | A*02:01 | YLQPRTFLL | TRAV12-1*0 | TRAJ30*01 | CVVNRDDKII | TRBV7-9*01 | TRBJ1-1*01 | CASGHPNDL | -1.551 |
| human | A*02:01 | YLQPRTFLL | TRAV12-1*0 | TRAJ31*01 | CVVNHLDRL | TRBV7-9*01 | TRBJ2-1*01 | CASSHDISLF | -0.401 |
| human | A*02:01 | YLQPRTFLL | TRAV12-1*0 | TRAJ31*01 | CVVNKEDRL | TRBV30*01 | TRBJ2-2*01 | CAFGQQNTG | -2.908 |
| human | A*02:01 | YLQPRTFLL | TRAV12-1*0 | TRAJ31*01 | CVVNKEDRL | TRBV5-1*01 | TRBJ2-2*01 | CASGDTNTG | -2.9 |
| human | A*02:01 | YLQPRTFLL | TRAV12-1*0 | TRAJ31*01 | CVVNKEDRL | TRBV7-8*01 | TRBJ2-2*01 | CASHSDRNT | -0.987 |
| human | A*02:01 | YLQPRTFLL | TRAV12-1*0 | TRAJ31*01 | CVVNVRDRL | TRBV7-9*01 | TRBJ1-1*01 | CASTDDIEAF | -0.259 |
| human | A*02:01 | YLQPRTFLL | TRAV12-1*0 | TRAJ34*01 | CVVNEADKLI | TRBV9*01 | TRBJ2-2*01 | CASSEQNTG | -3.601 |
| human | A*02:01 | YLQPRTFLL | TRAV12-1*0 | TRAJ34*01 | CVVNEEDKLI | TRBV7-9*01 | TRBJ2-2*01 | CAGSTSLTGE | -1.467 |
| human | A*02:01 | YLQPRTFLL | TRAV12-1*0 | TRAJ34*01 | CVVNEGDKLI | TRBV7-9*01 | TRBJ2-2*01 | CASANPDTG | -1.981 |
| human | A*02:01 | YLQPRTFLL | TRAV12-1*0 | TRAJ34*01 | CVVNGNTDK | TRBV7-9*01 | TRBJ2-1*01 | CASNEQNSN | -2.446 |
| human | A*02:01 | YLQPRTFLL | TRAV12-1*0 | TRAJ34*01 | CVVNKGDKLI | TRBV7-9*01 | TRBJ1-1*01 | CASSPDIEAFI | -0.106 |
| human | A*02:01 | YLQPRTFLL | TRAV12-1*0 | TRAJ34*01 | CVVNSFDKLI | TRBV7-9*01 | TRBJ2-7*01 | CASSLEIEQY | -0.821 |
| human | A*02:01 | YLQPRTFLL | TRAV12-1*0 | TRAJ34*01 | CVVNYDTDKI | TRBV2*01 | TRBJ2-2*01 | CATGGLNTG | -3.087 |
| human | A*02:01 | YLQPRTFLL | TRAV12-1*0 | TRAJ39*01 | CVVNNAGNI | TRBV15*01 | TRBJ2-2*01 | CATQNLNTG | -2.567 |

...

https://github.com/phbradley/TCRdock/blob/main/datasets_from_the_paper/table_S2_specificity_benchmark_tcrs.csv

**Supplementary Table 3: Decoy peptides for the specificity prediction benchmark**

| organism | mhc | peplen | peptide |
|---|---|---|---|
| human | A*02:01 | 9 | FLVFDSHYL |
| human | A*02:01 | 9 | GMYQGVADV |
| human | A*02:01 | 9 | ILIAWNPLM |
| human | A*02:01 | 9 | LLADAYRGV |
| human | A*02:01 | 9 | LTISGIYRV |
| human | A*02:01 | 9 | RLQTEIRNV |
| human | A*02:01 | 9 | SLWLPTASI |
| human | A*02:01 | 9 | TLILKLPSV |
| human | A*02:01 | 9 | YIDAPQVAI |
| human | A*02:01 | 10 | ALHYPKDIFL |
| human | A*02:01 | 10 | ALYPSLGSTI |
| human | A*02:01 | 10 | FLNWGTTKSL |
| human | A*02:01 | 10 | GLFPSQMKHA |
| human | A*02:01 | 10 | KLALPAAVSL |
| human | A*02:01 | 10 | NLSPGPVGYL |
| human | A*02:01 | 10 | SLALGLESQV |
| human | A*02:01 | 10 | SLWISSVERL |
| human | A*02:01 | 10 | TLTISGIYRV |
| mouse | H2Db | 9 | AIPRNILLL |
| mouse | H2Db | 9 | ASLPNVDIL |
| mouse | H2Db | 9 | KAYSVGHPM |
| mouse | H2Db | 9 | KSMSHMNSM |
| mouse | H2Db | 9 | SAIPMICGM |
| mouse | H2Db | 9 | STIRDGPAL |
| mouse | H2Db | 9 | TSGENAVSI |
| mouse | H2Db | 9 | VATTNLDYI |
| mouse | H2Db | 9 | YTTTNTLFI |
| mouse | H2Db | 10 | AASLPNVDIL |
| mouse | H2Db | 10 | ASLPNVDILT |
| mouse | H2Db | 10 | KSALNKATKL |
| mouse | H2Db | 10 | LAIPRNILLL |
| mouse | H2Db | 10 | QTEIRNVESF |
| mouse | H2Db | 10 | RLLLNFESCI |
| mouse | H2Db | 10 | TEIRNVESFL |
| mouse | H2Db | 10 | YLVPIKSDEM |
| mouse | H2Db | 10 | YQGVADVATL |

https://github.com/phbradley/TCRdock/blob/main/datasets_from_the_paper/table_S3_specificity_benchmark_decoy_peptides.csv

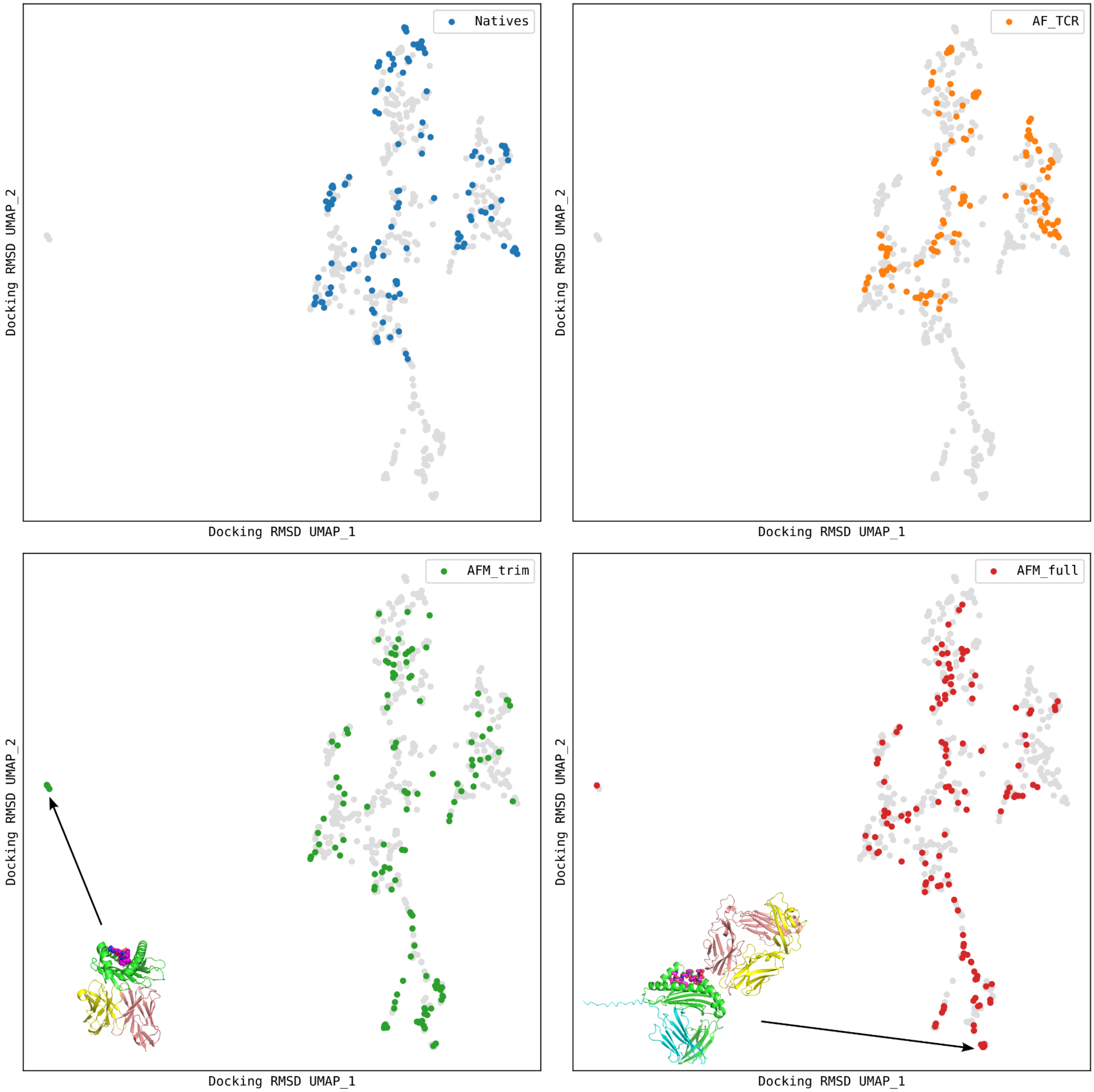


**Supplementary Figure 1. Docking geometry landscapes for the structure prediction benchmark.** The UMAP (McInnes et al., 2018) algorithm was used to transform a matrix of docking RMSD distances into a 2D landscape projection of the 130 native structures and the models from the AlphaFold TCR pipeline ('AF_TCR') and from two variants of the AlphaFold-Multimer model ('AFM_trim' and 'AFM_full'). Regions of docking space not sampled by the natives can be seen at the far left and bottom of the landscapes. Representative AlphaFold-Multimer models are shown: in the bottom left panel, a TCR docked to the underside of the MHC; in the bottom right panel, a TCR docked to the side of the MHC and not making contacts with the peptide.

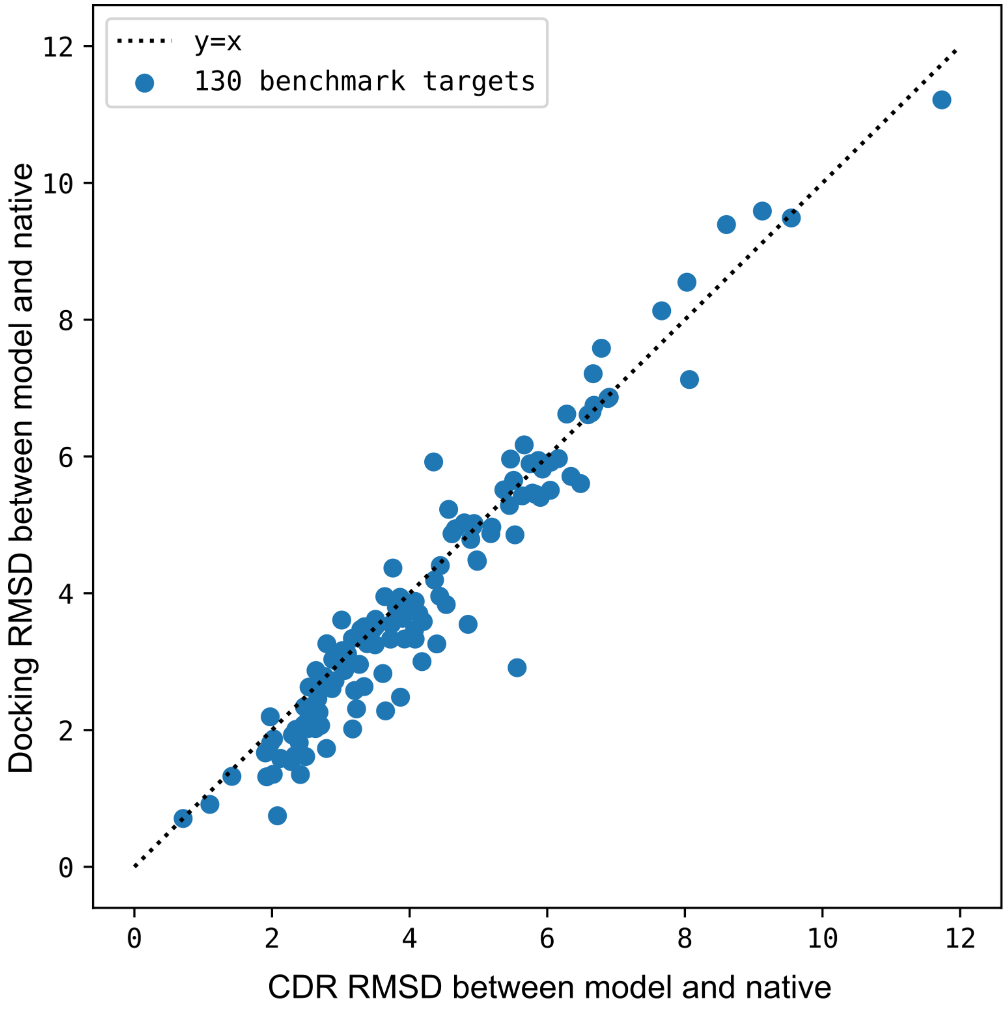


**Supplementary Figure 2. Comparison of docking RMSD to CDR RMSD.** Docking RMSD and CDR RMSD are well correlated. CDR RMSD tends to be larger than docking RMSD because it also includes deviation in the internal structures of the CDR loops.

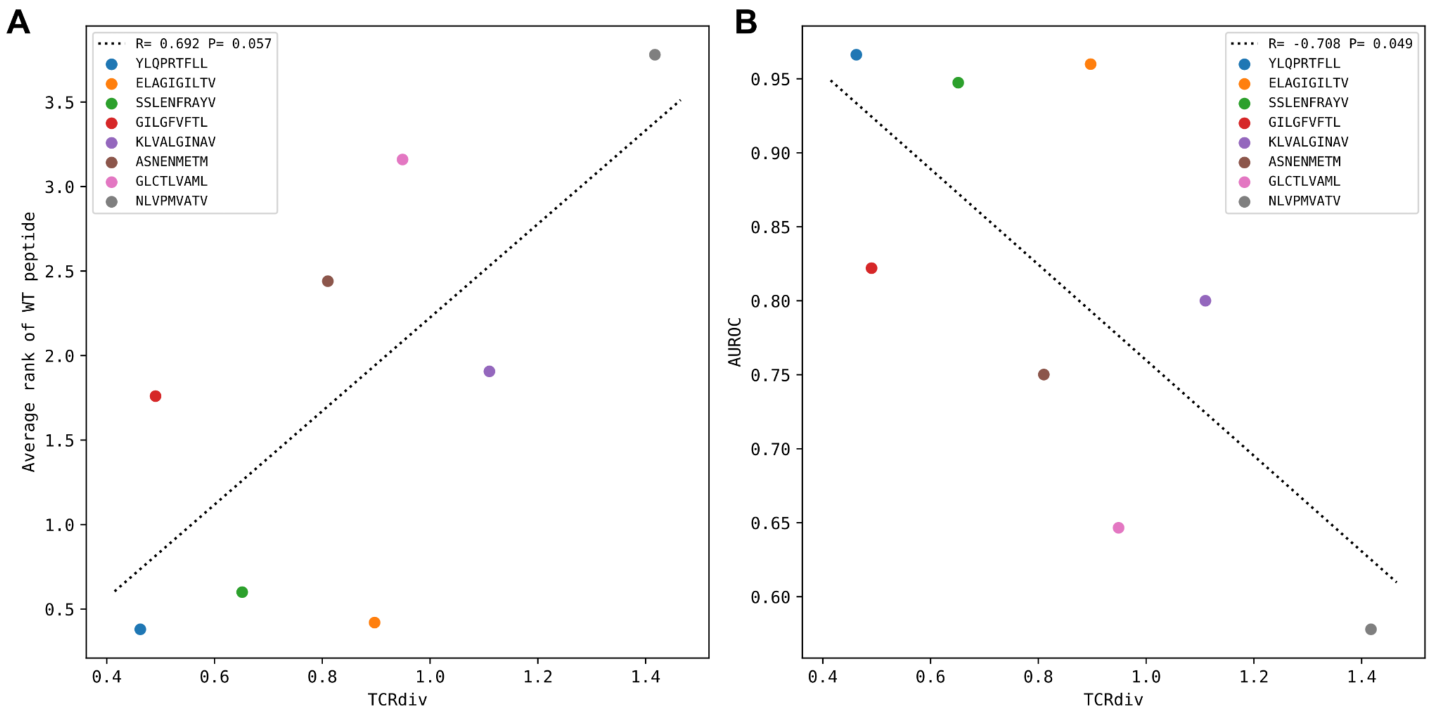


**Supplementary Figure 3. Peptide specificity prediction accuracy is inversely correlated with repertoire sequence diversity.** TCR repertoire sequence diversity is measured with the TCRdiv measure (Dash et al., 2017) with a sigma value of 120. Each colored marker corresponds to one of the eight benchmark epitopes as indicated in the legend. **(A)** For each epitope, the average rank of the wild type peptide relative to the nine decoys (0=best binding score; 9=worst binding score) is calculated over all the epitope-specific TCRs and plotted against the TCRdiv score for the corresponding TCR repertoire. **(B)** TCRdiv scores are plotted against the area under the receiver operating characteristic curve (AUROC) values for discriminating the wild type from decoy peptides by binding score.

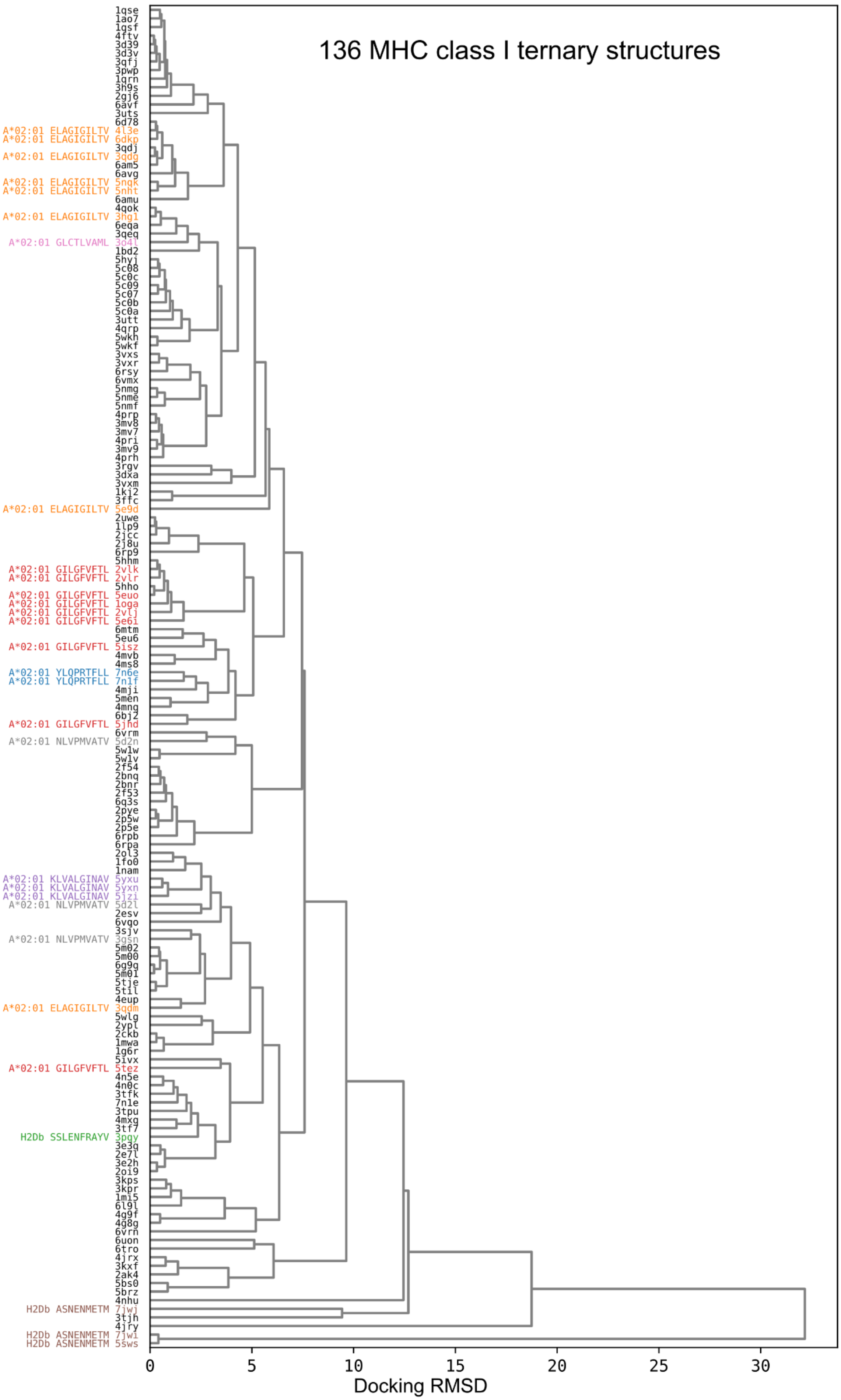

136 MHC class I ternary structures
1qse
1ao7
1qsf
4ftv
3d39
3d3v
3qfj
3pwp
1qrn
3h9s
2gj6
6avf
3uts
6d78
A*02:01 ELAGIGILTV 4l3e
A*02:01 ELAGIGILTV 6dkp
3qdj
A*02:01 ELAGIGILTV 3qdg
6am5
6avg
A*02:01 ELAGIGILTV 5nqk
A*02:01 ELAGIGILTV 5nht
6amu
4qok
A*02:01 ELAGIGILTV 3hg1
6eqa
3qeq
A*02:01 GLCTLVAML 3o4l
1bd2
5hyj
5c08
5c0c
5c09
5c07
5c0b
5c0a
3utt
4qrp
5wkh
5wkf
3vxs
3vxr
6rsy
6vmx
5nmg
5nme
5nmf
4prp
3mv8
3mv7
4pri
3mv9
4prh
3rgv
3dxa
3vxm
1kj2
3ffc
A*02:01 ELAGIGILTV 5e9d
2uwe
1lp9
2jcc
2j8u
6rp9
5hhm
A*02:01 GILGFVFTL 2vlk
A*02:01 GILGFVFTL 2vlr
5hho
A*02:01 GILGFVFTL 5euo
A*02:01 GILGFVFTL 1oga
A*02:01 GILGFVFTL 2vlj
A*02:01 GILGFVFTL 5e6i
6mtm
5eu6
A*02:01 GILGFVFTL 5isz
4mvb
4ms8
A*02:01 YLQPRTFLL 7n6e
A*02:01 YLQPRTFLL 7n1f
4mji
5men
4mng
6bj2
A*02:01 GILGFVFTL 5jhd
6vrm
A*02:01 NLVPMVATV 5d2n
5w1w
5w1v
2f54
2bnq
2bnr
2f53
6q3s
2pye
2p5w
2p5e
6rpb
6rpa
2ol3
1fo0
1nam
A*02:01 KLVALGINAV 5yxu
A*02:01 KLVALGINAV 5yxn
A*02:01 KLVALGINAV 5jzi
A*02:01 NLVPMVATV 5d2l
2esv
6vqo
3sjv
A*02:01 NLVPMVATV 3gsn
5m02
5m00
6g9q
5m01
5tje
5til
4eup
A*02:01 ELAGIGILTV 3qdm
5wlg
2ypl
2ckb
1mwa
1g6r
5ivx
A*02:01 GILGFVFTL 5tez
4n5e
4n0c
3tfk
7n1e
3tpu
4mxg
3tf7
H2Db SSLENFRAYV 3pqy
3e3q
2e7l
3e2h
2oi9
3kps
3kpr
1mi5
6l9l
4g9f
4g8g
6vrn
6uon
6tro
4jrx
3kxf
2ak4
5bs0
5brz
4nhu
H2Db ASNENMETM 7jwj
3tjh
4jry
H2Db ASNENMETM 7jwi
H2Db ASNENMETM 5sws
0
5
10
15
20
25
30
Docking RMSD

**Supplementary Figure 4. Hierarchical clustering tree of TCR:pMHC class I docking geometries.** Structures are labeled with PDB IDs and, for the epitopes in the peptide decoy discrimination benchmark, MHC and peptide sequence colored as in main text **Figures 3, 5,** and **6**.

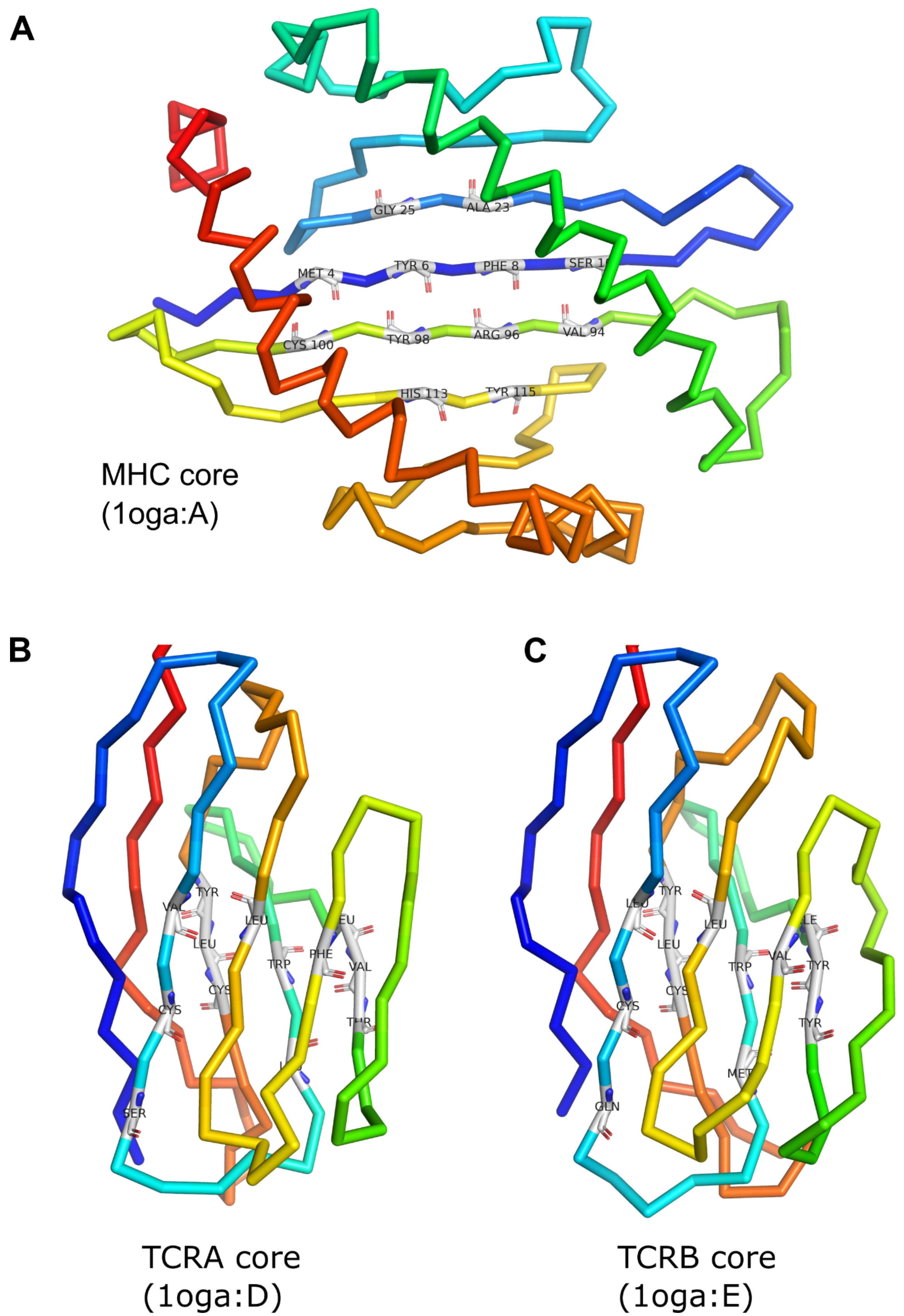


**Supplementary Figure 5. MHC and TCR core residue definitions.** The MHC symmetry axis is the rotation axis of the transformation that superimposes the 12 core residues onto themselves, interchanging the N- and C-terminal half-cores. The TCR symmetry axis is the rotation axis of the superposition transform that interchanges the TCRA and TCRB cores.

##### Supplementary Algorithm 1. TCRdiv calculation implemented in Python.

```
import numpy as np
def compute_tcrdiv(D, sigma=120.):
    ''' D is a symmetric matrix of TCRdist distances
    sigma is the width of the Gaussian smoothing term
    '''
    N = D.shape[0]
    D[np.arange(N), np.arange(N)] = 1e6 # set diagonal to a very large value
    return -1*np.log(np.sum(np.exp(-1*(D/sdev)**2))/(N*(N-1)))
```

##### Supplementary Algorithm 2. Kernel-density based algorithm for subsampling implemented in Python

```
import numpy as np
def pick_reps(D, num_reps=50, sdev_big=120., sdev_small=36., min_size=0.5):
    ''' D is a symmetric distance matrix (e.g., of TCRdist distances)
    num_reps is the number of representatives to choose
    sdev_big defines the neighbor-density sum used for ranking
    sdev_small limits the redundancy
    both sdev_big and sdev_small are in distance units (ie same units as D)
    '''
    # the weight remaining for each instance
    wts = np.array([1.0]*D.shape[0])

    reps, sizes = [], []
    for ii in range(num_reps):
        if np.sum(wts)<1e-2:
            break
        gauss_big   = np.exp(-1*(D/sdev_big  )**2) * wts[:,None] * wts[None,:]
        gauss_small = np.exp(-1*(D/sdev_small)**2) * wts[:,None] * wts[None,:]
        nbr_sum = np.sum(gauss_big, axis=1)
        rep = np.argmax(nbr_sum)
        size = nbr_sum[rep]
        if size<min_size:
            break
        wts = np.maximum(0.0, wts - gauss_small[rep,:]/wts[rep])
        assert wts[rep] < 1e-3
        reps.append(rep)
        sizes.append(size)
    return reps, sizes
```
